# Rapid and highly-specific generation of targeted DNA sequencing libraries enabled by linking capture probes with universal primers

**DOI:** 10.1101/422519

**Authors:** Joel Pel, Amy Leung, Wendy W. Y. Choi, Milenko Despotovic, W. Lloyd Ung, Gosuke Shibahara, Laura Gelinas, Andre Marziali

## Abstract

Targeted Next Generation Sequencing (NGS) is being adopted increasingly broadly in many research, commercial and clinical settings. Currently used target capture methods, however, typically require complex and lengthy (sometimes multi-day) workflows that complicates their use in certain applications. In addition, small panels for high sequencing depth applications such as liquid biopsy typically have low on-target rates, resulting in unnecessarily high sequencing cost.

We have developed a novel targeted sequencing library preparation method, named Linked Target Capture (LTC), which replaces typical multi-day target capture workflows with a single-day, combined ‘target-capture-PCR’ workflow. This approach uses physically linked capture probes and PCR primers and is expected to work with panel sizes from 100 bp to >10 Mbp. It reduces the time and complexity of the capture workflow, eliminates long hybridization and wash steps and enables rapid library construction and target capture. High on-target read fractions are achievable due to repeated sequence selection in the target-capture-PCR step, thus lowering sequencing cost.

We have demonstrated this technology on sample types including cell-free DNA (cfDNA) and formalin-fixed, paraffin-embedded (FFPE) derived DNA, capturing a 35-gene pan-cancer panel, and therein detecting single nucleotide variants, copy number variants, insertions, deletions and gene fusions. With the integration of unique molecular identifiers (UMIs), variants as low as 0.25% abundance were detected, limited by input mass and sequencing depth. Additionally, sequencing libraries were prepared in less than eight hours from extracted DNA to loaded sequencer, demonstrating that LTC holds promise as a broadly applicable tool for rapid, cost-effective and high performance targeted sequencing.

## Introduction

Targeted Next Generation Sequencing (NGS) is common practice in many research, commercial and clinical applications, as a faster and cheaper alternative to equivalent depth whole-genome or whole-exome sequencing. As sequencing technologies continue to become more accessible, the adoption of targeted NGS into more labs and markets is likely to follow.

Existing targeted sequencing approaches generally fall into three categories: (i) Multiplexed PCR; (ii) Hybridization and extension; and (iii) Hybridization and capture (1), and are summarized briefly here. PCR is a well-known technique which can be very effective for targeting small to mid-sized genomic regions. However, multiplex PCR is generally challenging to design and does not scale easily to very large targets. Sample splitting is generally required to tile large contiguous regions or reduce primer dimers, subsequently reducing sensitivity to rare variants (2). Techniques aimed at mitigating multiplexing challenges include using droplets to reduce primer dimer formation (3), integrating special primer adapters to enable tiling without sample splitting (4), or linking primers to increase specificity and reduce primer dimers (5, 6). While providing improvements, these methods are generally more complex to design and use, and are still limited in their multiplexing capabilities. Additionally, for many applications, including diagnostics, PCR methods generally lose information compared to ligation-based methods. For example, in multiplex PCR methods, the start and stop positions of genomic fragments are lost, and integration of unique molecular identifiers (UMIs) for somatic mutation detection can be challenging (7).

Hybridization and extension methods improve on PCR multiplexing limitations by using a single ‘primer’ for each target that extends across a region of interest and reduces primer dimers (8-12). The resulting products are then ligated and amplified by universal primers to create sufficient material for sequencing. Despite the improvements in multiplexing compared to PCR due to fewer primers, these methods have not achieved the same widespread use compared to hybridization and capture methods. Potential reasons may include high DNA input mass requirements, high cost and complexity, low uniformity, or loss of sequence information under long priming regions (1, 4).

Perhaps the most common approach, hybridization and capture (13, 14), uses single-stranded DNA or RNA probes that are designed to bind specifically to sequences of interest. Probes containing biotin are annealed to targets during a lengthy incubation step, after which avidin-biotin binding is used to extract the biotin-labeled probes, thus enriching for the targets of interest. Hybridization and capture methods have many advantages, including scalability to large panels, the ability to easily distinguish duplicates on the sequencer through use of UMIs, and to retain insert start-stop positions due to up-front ligation. Some of the main disadvantages, however, include low sequencer on-target fraction, high cost, and complex and lengthy workflows (4).

Commercial hybridization and capture methods vary in speed, complexity and performance. These methods typically start with a library preparation step (either by ligation or transposase), followed by a universal pre-amplification PCR step and then one or more hybridization capture steps, ranging from four to 72 hours. Following capture, the targeted DNA is recovered via a series of pull-down and wash steps. Targeted DNA is then amplified again and quantified prior to sequencing (15). In general, faster capture times can only be achieved at the expense of lower on-target fractions. Also, as panel size decreases from ∼30 Mbp for whole exome captures to the 10 kbp −100 kbp range commonly used for diagnostic applications, on-target fraction generally decreases as well (16). Lower on-target results in lower depth of coverage and lower variant sensitivity unless sequencing throughput (and cost) is increased (15, 17).

To the best of our knowledge, the IDT xGen workflow (Integrated DNA Technologies) is the fastest available commercial assay, with a reported workflow time of nine hours. However, this does not include library preparation or pre-amplification which generally adds at least several more hours (depending on method), requiring the workflow to be performed over multiple work days. Other common protocols can span two or more days, such as Roche SeqCap (Roche Sequencing). The length and complexity of these workflows limit their use, especially in clinical settings, where fast turn-around time and ease of use are important.

We have developed Linked Target Capture (LTC), a novel target capture method with broad application, designed to reduce hybridization workflows to less than eight hours while retaining high performance over all panel sizes. LTC replaces existing hybridization methods with a combined ‘target-capture-PCR’ workflow using linked capture probes and universal amplification primers. Here we describe the LTC method, and demonstrate its ability to rapidly deliver enriched sequencing libraries from multiple sample types, including formalin-fixed, paraffin-embedded (FFPE) derived DNA, plasma-derived cell-free DNA (cfDNA) and cell line DNA. Additionally, with the integration of UMIs, we demonstrate LTC’s ability to detect low-level single nucleotide variants, copy number variants, insertions/deletions and gene fusions.

## Results

### Linked Target Capture Concept

The LTC method is illustrated in Figure 1 for Illumina sequencers, though it is expected to be compatible with most sequencing platforms. The workflow begins with ligation of short Y-adapters that contain truncated portions of the Illumina P5 and P7 flow cell binding sequences, such that ligated molecules will not bind to the flow cell and be sequenced without further processing. Following ligation and pre-amplification using universal primers, two sequential target-capture-PCR (tcPCR) steps are performed with Probe-Dependent-Primers (PDPs). PDPs consist of non-extendable DNA capture probes linked 5’ to 5’ with a low melting-temperature universal primer complementary to a portion of the ligated adapter (Figure 1 (ii) and (iii)). When bound to their targets, the probes bring the universal primer into close proximity with the universal priming site on the template, increasing the reaction rate of primer binding and initiating polymerase extension. The polymerase displaces or digests the probe portion of the PDP to make a copy of the entire target template, and the reaction proceeds to the next tcPCR cycle. To create sequencer-compatible libraries, the second tcPCR integrates the full Illumina P5 and P7 sequences into the universal primer portion of the PDPs. Both tcPCR reactions are performed above the melting temperature of the universal primers so that amplification is heavily biased towards target-bound PDPs.

**Figure 1:**
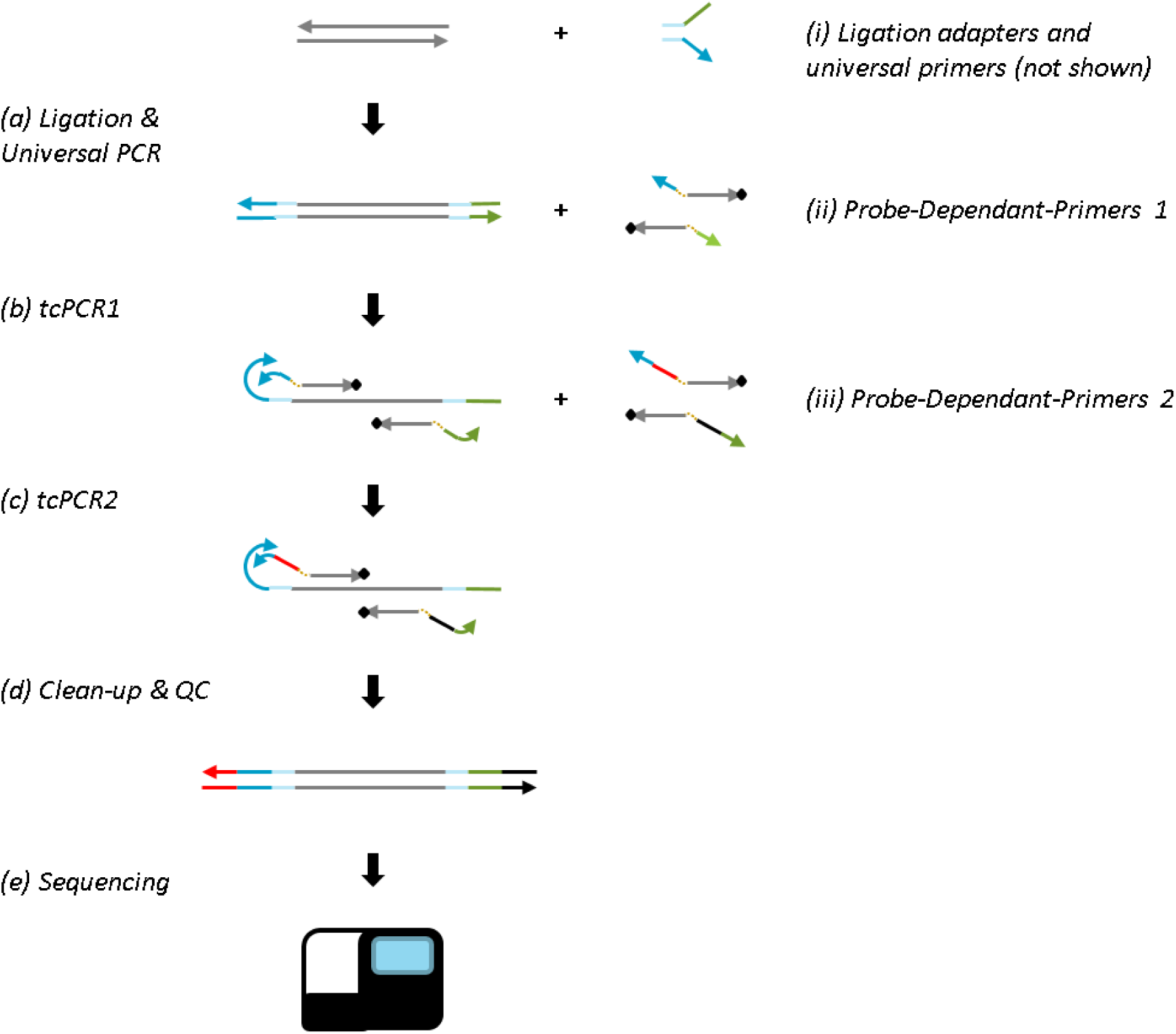
Linked Target Capture Workflow. (a) Custom adapters (i) are ligated to template DNA and the resulting product is amplified with universal primers. (b) Target regions are selectively amplified using custom probe-dependent-primers (PDPs) (ii) which contain a recognition sequence (dark grey) with a 3’ blocker (black diamond) and are linked to an oligo containing a universal priming sequence for the first target capture PCR reaction (tcPCR1). (c) A second set of PDPs (iii), which contain Illumina adapters (red and black) between the probe and linked universal primer, are then added and a second target capture PCR reaction (tcPCR2) is completed prior to (d), clean up and QC and (e) loading on a sequencer.

As described in the Materials and Methods, PDPs are made by reacting separately synthesized probes and primers. PDP panels are made by linking probe sets to the universal primers, making panel generation, expansion, and combination straightforward.

### Assay Validation

To validate the LTC workflow, PDPs were designed to cover relevant portions of 35 cancer-related genes, as described in Materials and Methods and listed in S1 Table. PDPs were chosen to capture four major variant types, including single nucleotide variants (SNVs), insertions/deletions (Indels), copy number variants (CNVs) and structural rearrangements (ex: gene fusions). Libraries were created and sequenced in duplicate from four sample types, as outlined in Table 1: mechanically sheared cell line DNA, enzymatically sheared cell line DNA, cfDNA, and FFPE-derived DNA. Additionally, to test lower input mass, duplicate libraries were created and sequenced from 5 ng of mechanically sheared cell line DNA. The total time from extracted DNA to loaded sequencer was eight hours, with about three hours of hands-on time.

**Table 1:** LTC 35-gene sequencing performance data for two replicates of sample type and DNA mass. On-target fraction was defined as the fraction of total bases that aligned to the target regions. Mean target coverage was defined as the mean de-duplicated coverage over all target regions, and uniformity was defined as the fraction of on-target bases that were covered within two fold of the mean target coverage (i.e. between 0.5x and 2x of the mean). Median insert length was measured over all de-duplicated on-target inserts.

| DNA Source | DNA Mass | Replicate | On-target Fraction | Mean Target Coverage | Uniformity | Median Insert Length |
| --- | --- | --- | --- | --- | --- | --- |
| **Mechanically sheared cell line** | **5** | 1 | 93% | 157 | 95% | 162 |
|  |  | 2 | 93% | 156 | 95% | 162 |
|  | **50** | 1 | 93% | 1202 | 96% | 168 |
|  |  | 2 | 89% | 1281 | 96% | 169 |
| **Enzymatically sheared cell line** | **50** | 1 | 91% | 1685 | 94% | 193 |
|  |  | 2 | 86% | 1764 | 93% | 190 |
| **FFPE tissue** | **40** | 1 | 91% | 1358 | 96% | 175 |
|  |  | 2 | 86% | 1444 | 96% | 171 |
| **Plasma cfDNA** | **50** | 1 | 95% | 1154 | 90% | 174 |
|  |  | 2 | 97% | 1148 | 90% | 175 |

All libraries were analyzed through the same pipeline (see Materials and Methods) and down-sampled to a fixed number of sequencing clusters (or read pairs) for a given input mass (2 M read pairs for 40 or 50 ng, 0.2 M for 5 ng). Fixing the number of read pairs is important when comparing results, as the same sequencing data analyzed with different numbers of read pairs produces different results (especially in coverage). This is attributed to several factors, including insufficient reads for a given input mass (or a given number of input genomes), and Poisson variation. Fixed-read results are shown in Table 1. On-target fraction, mean target coverage and uniformity were calculated using Picard CollectHSMetrics (broadinstitute.github.io/picard/), as described in Materials and Methods.

These data demonstrate consistently high on-target fraction (86%-97%) and uniformity (90%-96%) across a range of sample types and input mass relevant to clinical applications of targeted sequencing. As a reference, commercially available Roche, Illumina and Agilent methods have been compared using a 110 gene panel, and ranged in performance from 75% to ∼87% on-target (17). While not a direct comparison, this reference provides a good indicator of relative performance, as it is typically easier to achieve high on-target fraction on large panels (16) (a direct in-house comparison was not undertaken due to the significant cost of capture panels). To demonstrate the scalability of LTC, we measured enrichment on four of the 35 gene targets (BRAF, EGFR, ERBB2 and TP53), using 50 ng mechanically-sheared cell line DNA. The measured on-target fraction was >97% in both replicates, higher than the same measurement for our 35-gene panel. Similarly-sized small panels using conventional single-round target capture reported ∼5% on-target reads in (16) and (18).

A comparison in uniformity can be made against the use of a SureSelect XT panel (Agilent Technologies) covering 231 SNV targets in 26 genes (19). For FFPE samples with similar coverage (>1000x) in (19), the authors report uniformity ranging from ∼50% to 93%, whereas both FFPE replicates using LTC had a uniformity of 96%.

Insert length distributions for each sample type were calculated using Picard CollectInsertSizeMetrics and are shown in Figure 2. Mechanically sheared cell lines were created by the manufacturer to produce a majority of inserts in the range of ∼100 bp to 250 bp (see Materials and Methods). The recovered insert lengths for these samples represent a good match to the expected size distribution with 89% of targets between 100 bp and 250 bp. Enzymatically-sheared DNA samples produced slightly longer inserts, likely a function of the shearing protocol. Additionally, the median insert size for the cfDNA samples was 175 bp in a reasonably narrow distribution, in good concordance with literature (20). A peak was also visible around ∼325 bp, suggesting these long fragments may have been wrapped twice around the histone. Finally, FFPE-derived DNA samples also produced a short insert length distribution, as expected from the degradation associated with FFPE samples combined with enzymatic shearing and repair.

**Figure 2:**
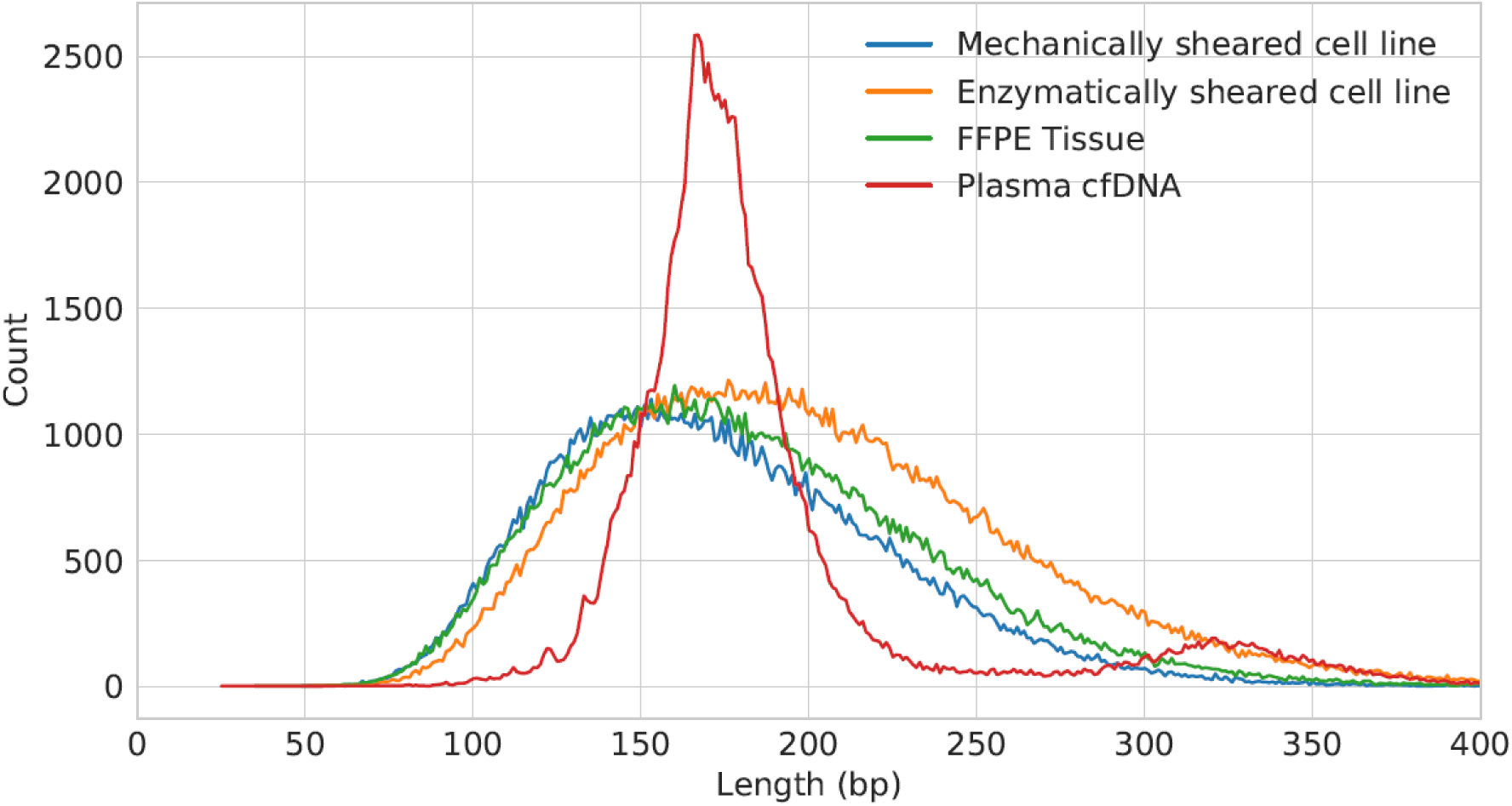
Representative insert size distributions for each sample type used in this study.

### Variant Detection

To enable the detection of low level variants with LTC, UMIs consisting of four random bases in series were integrated into Illumina’s ‘Index 1’ read position of the ligation adapter. The UMIs were used in conjunction with the start and stop positions of the inserts to uniquely identify individual starting template molecules and to create consensus sequences (see Materials and Methods). A commercially available reference standard cell line (HD786, Horizon Discovery) was used to assess the ability of LTC to detect variants as it contains SNVs, CNVs, indels and fusions at levels characterized by the manufacturer. The variants covered by the 35-gene panel are listed in Table 2, along with the expected allele percentage as specified by the manufacturer for each of the different samples used in this study.

**Table 2:**
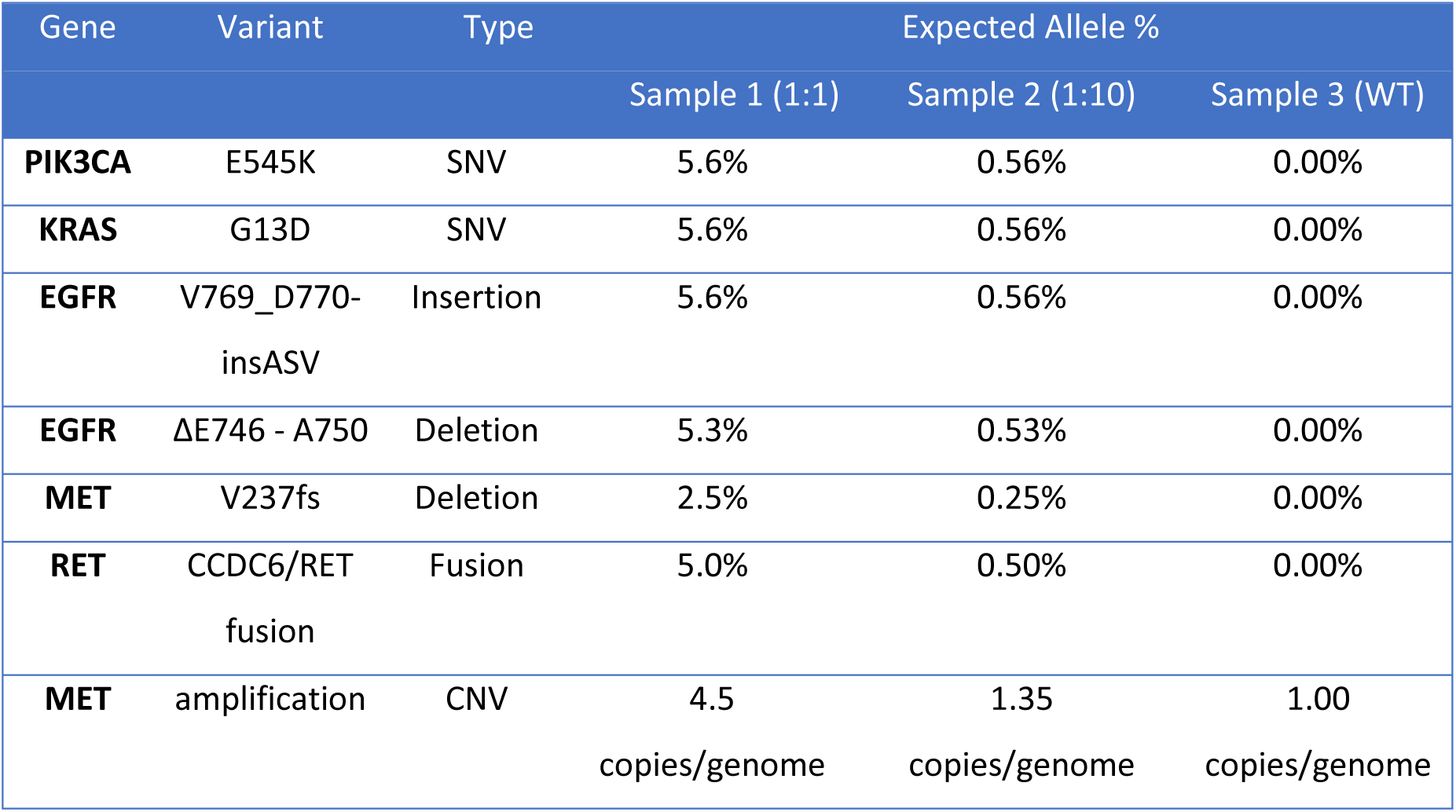
Reference standard variants. The expected allele percentage was measured and specified by the manufacturer using digital PCR or next generation sequencing. Expected allele percentages are given for stock samples (Sample 1), samples diluted to 1/10 of the stock concentration (Sample 2), and wild-type samples (Sample 3).

| Gene | Variant | Type | Expected Allele % |  |  |
| --- | --- | --- | --- | --- | --- |
|  |  |  | Sample 1 (1:1) | Sample 2 (1:10) | Sample 3 (WT) |
| **PIK3CA** | E545K | SNV | 5.6% | 0.56% | 0.00% |
| **KRAS** | G13D | SNV | 5.6% | 0.56% | 0.00% |
| **EGFR** | V769_D770-insASV | Insertion | 5.6% | 0.56% | 0.00% |
| **EGFR** | ΔE746 - A750 | Deletion | 5.3% | 0.53% | 0.00% |
| **MET** | V237fs | Deletion | 2.5% | 0.25% | 0.00% |
| **RET** | CCDC6/RET fusion | Fusion | 5.0% | 0.50% | 0.00% |
| **MET** | amplification | CNV | 4.5<br>copies/genome | 1.35<br>copies/genome | 1.00<br>copies/genome |

To test variant detection, 50 ng of DNA was used from each cell line. DNA from the reference standard (Sample 1) was analyzed in duplicate, along with duplicate analysis of cell line from the same manufacturer known to be wild type for the variants of interest (Sample 3). A ten-fold titration (Sample 2) of the reference standard was made with the wild type cell line, and also tested in duplicate. Sequencing analysis and variant calling was performed as outlined in Materials and Methods.

The measured variant fractions for detected SNVs, indels and fusions are plotted against the expected fractions in Figure 3. All the variants that were detected were measured at allele frequencies within ∼3x of expected values. Expected variants as low as 0.25% were detected (the lowest fraction tested in this study), which corresponds to ∼38 mutant fragments present in the initial 50 ng sample (assuming 3.3 pg of DNA per human haploid genome). Since ligation yield in general is much lower than 100% (21), the actual number of mutants entering capture could be considerably less than 38, and perhaps near sampling error for some loci. Discrepancy between measured and expected values may be attributed to a number of factors including the differences in variant calling methods, titration of the reference standard, and the relative sequencing coverage of each variant, all of which could potentially lead to sampling error.

**Figure 3:**
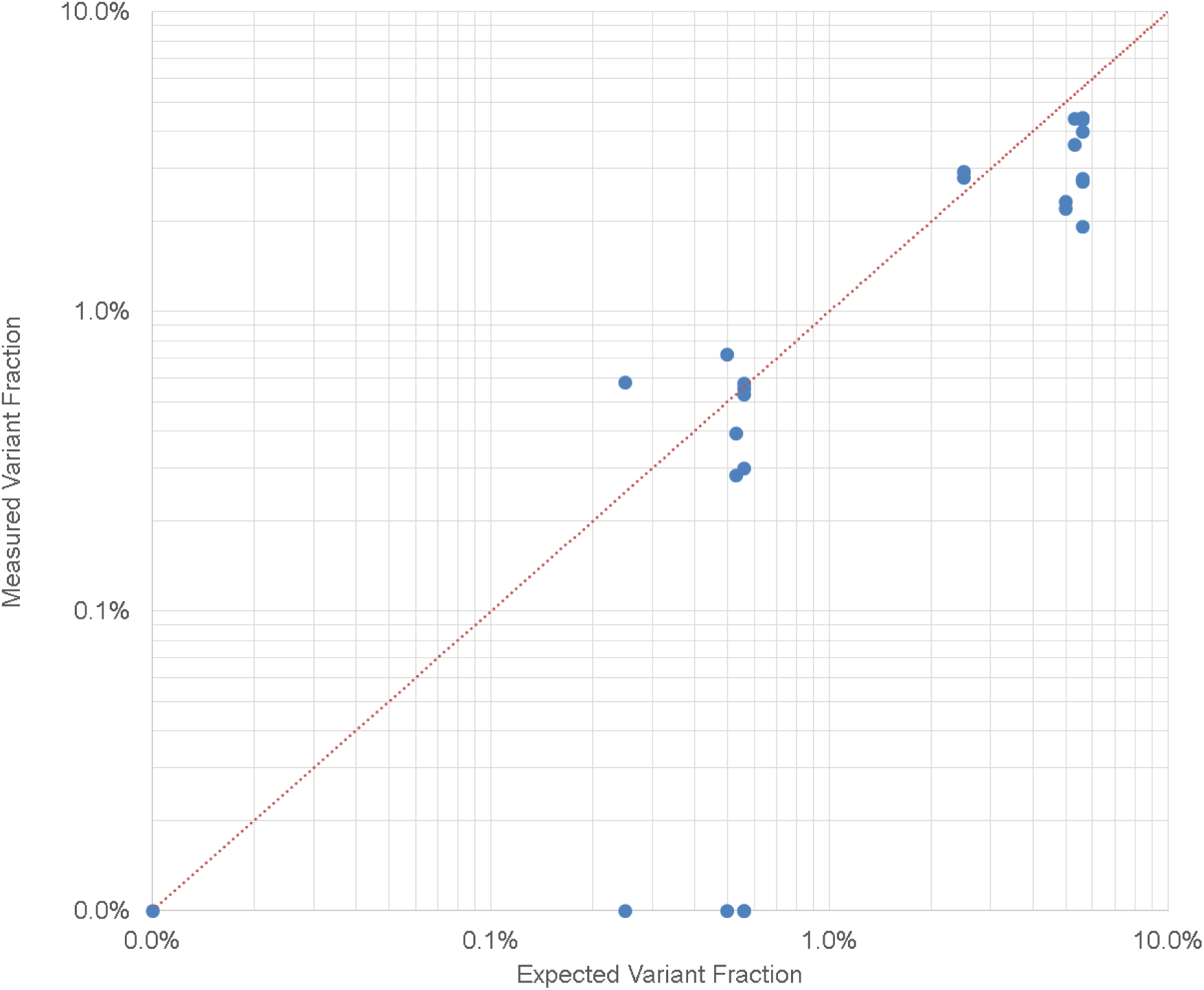
Expected vs. measured SNV, indel and fusion fractions. The dotted line represents a 1:1 ratio of expected vs. measured variants. Undetected and zero variant fraction samples were reported at 0.0% for display purposes.

All twelve (100%) of the SNV, indel and fusion variants were correctly identified at 0.00% variant fraction in the wild-type cell line (Sample 3 replicates). Eight of the twelve (67%) SNV, indel and fusion variants expected between 0.25% and 0.56% were detected in the diluted reference standard (Sample 2 replicates), while all twelve (100%) of the same variants expected between 2.5% and 5.6% were detected in the reference standard (Sample 1 replicates). With further improvements to LTC (see Discussion), we expect even higher sensitivity and lower detection limits to be possible.

Copy number variation was assessed for the MET gene in all six samples (replicates of Samples 1, 2 and 3) by our analysis pipeline, which was designed to identify samples as “amplified”, “deleted” or “copy-number neutral” (see Materials and Methods). The MET gene was identified as “amplified” in both replicates of Sample 1, and “copy-number neutral” for both replicates of Samples 2 and 3. These results were consistent with expectations, as the 4.5 copies of the MET gene present in each Sample 1 replicate should be easily detectable above background, even when compared against only two wild-type samples (Sample 3 replicates). On the other hand, the 1.35 copies of the MET gene in each of the Sample 2 replicates would likely require many more measurements to confidently detect a copy number variation above the wild-type background. In general, all four variant types were detected as expected, demonstrating the capability of LTC as a target capture tool for many different applications.

## Discussion

In general, target capture performance and workflow improvements have the potential to increase NGS and target capture usage in existing applications, and also to enable new opportunities if workflow time, complexity or cost are reduced.

Arguably the most significant improvement of LTC over existing methods is the dramatic decrease in workflow time. To the best of our knowledge, the IDT xGen workflow is the fastest commercial hybridization capture method, at nine hours. This does not include library preparation, which generally adds several additional hours, and requires the assay to be run over two work days. In contrast, the LTC workflow was completed in eight hours, including library preparation and loading of the sequencer.

Typical capture workflows are limited by the length and performance of the hybridization step, which on its own can extend to 72 hours. Shorter hybridization steps typically compromise performance resulting in either lower de-duplicated target coverage or higher off-target fraction. LTC avoids this tradeoff and shortens this rate limiting step by employing a combined target-capture-PCR (tcPCR) step. High de-duplicated target coverage is achieved by specifically capturing both senses (see Materials and Methods) and by operating at a relatively low temperature compared to the probe binding temperature, while the on-target fraction is increased through many effective capture cycles performed in each tcPCR reaction. An additional benefit of tcPCR is workflow simplicity. In conventional target capture workflows, biotinylated probes typically require binding to streptavidin coated beads to enrich for the target DNA. The subsequent bead capture and wash steps are generally complicated, labor intensive and can be difficult to automate (22), potentially limiting deployment of target capture workflows in some cases. On the other hand, the LTC tcPCR setup and operation are analogous to a standard PCR reaction, and thus are more familiar to a larger number of technicians, and also easier to automate. Additionally, it should be noted that LTC could be paired with any library preparation method that introduces the correct adapter sequences, such as single stranded library prep (23) or transposition (24).

A related advantage of the combined PCR-capture step is the ability to produce consistent sequencing performance from low input mass samples. Sequencing parameters, including coverage, scaled as expected from the 50 ng samples down to 5 ng, suggesting that LTC is able to recover molecules efficiently across a wide mass range. This is especially important in applications where sample is limiting, and could be tested to even lower limits in a future study.

It should be noted that the workflow time and complexity of LTC is comparable to multiplexed PCR (such as AmpliSeq by ThermoFisher) and hybridization extension methods (such as (11)). LTC holds a number of significant advantages over these methods, however. First, since the LTC primers are universal, it does not require sample splitting before amplification to prevent unwanted amplicon formation. This avoids loss of sensitivity and the requirement for large DNA input mass. Second, since LTC probes are displaced by the extended universal primer, sequence information at probe binding sites is retained on the amplified molecules to be sequenced, thus capturing all of the sequence information available from a single fragment. This is in contrast to PCR and hybridization extension methods where any variants contained within a PCR primer binding site are lost after the primer has bound and extended. LTC also retains fragment start and stop positions, which are lost in PCR and hybridization extension methods, and have been shown to provide useful biological information (25). Additionally, it is generally much easier to integrate low variant detection in hybridization capture methods like LTC compared to PCR methods. When UMIs are integrated in ligation as they are in LTC, it is easier to avoid labelling a single molecule with multiple UMIs, which can occur in PCR methods. Also, to our knowledge, it is not possible to integrate duplex sequencing in a PCR-based UMI method, but this has been demonstrated with LTC. Finally, because the challenges associated with multiplex PCR are reduced through the use of universal primers, the LTC workflow can be used for a wide range of panel sizes, including large panels for which multiplex PCR methods would not work. Small panels have been demonstrated in this study, and initial work towards larger panels indicates that exome-scale LTC panels may be possible. This is advantageous, as a single workflow could be implemented for multiple assays or applications.

LTC has several other unique properties. Primers and probes can be oriented to capture a specific strand of the target duplex DNA (ex: the sense strand, see Materials and Methods), providing an advantage in rare variant detection, or in applications where it is desirable to sequence only one strand of the starting template such as transcriptome sequencing (26). In addition, LTC has been demonstrated in droplets, providing multiplexing capabilities to droplet-based assays not achievable with standard capture methods.

The sequencing statistics achieved using Linked Target Capture were excellent, with greater than 91% average on-target and 94% average uniformity, providing cost-effective sequencer usage and leaving little room to improve these metrics. Measuring how these factors scale to much larger panels would be an important part of a future study. Mean target coverage was lower than initially expected, by about two to three fold compared to hybridization capture with similar analysis (27); we suspect this to be due to the lack of LTC probe tiling. The 35-gene panel used in this work consisted of fairly sparse probe placement to reduce panel cost, such that the probes covered less than 100% of bases in the targets. Initial data from tiling two targets in the 35-gene panel to nearly 200% demonstrated a more than 3-fold increase in mean target coverage, which agrees with previously reported tiling improvements of at least two-fold (28). It is expected that tiling will significantly improve mean target coverage as well as variant detection when applied across the whole 35-gene panel.

Variant detection may be further improved through the use of lower error UMI designs. Like hybridization and capture methods, the error rate of LTC is expected to be linked to the UMI design used in a given assay. For example, integrating duplex UMIs into the LTC ligation adapters is expected to further reduce the detection limit, similarly to the reduction observed for duplex UMIs applied to hybridization and capture methods (27). Increasing the input mass and sequencing depth are also expected to lower the detection limit of LTC.

In summary, we have developed a novel target capture method with a rapid workflow and efficient sequencer usage. With continued improvements in tiling and panel expansion, we expect LTC to be a high performance target capture method applicable in many settings.

## Materials and Methods

### PDP Design and Conjugation

In order to enable panel design flexibility, PDPs were made by conjugating target-specific probes and universal primers. The universal primers (forward and reverse) were manufactured by Integrated DNA Technologies (IDT) and contained a 5’ Dibenzocyclooctyl (DBCO) modification. The forward and reverse untailed primers for the first target capture step were CACCGAGATCT and TACGAGATCGG respectively. The forward and reverse tailed primers for the second target capture step were AATGATACGGCGACCACCGAGATCT and CAAGCAGAAGACGGCATACGAGATCGG respectively.

Capture probes were designed to cover portions of 35 cancer-related genes, shown in S1 Table. Total sequence coverage was 11,473 bp. Probes were designed with adjacent forward and reverse probes covering the desired regions, with zero gap between forward and reverse probes, a minimum length of 30 bp, maximum length of 70 bp, and a melting temperature of ∼85 °C calculated using uMELT (29) with default conditions. Probes were synthesized by IDT with a 5’ azide modification to conjugate with the DBCO on the primer and a 3’ inverted dT base, to inhibit polymerase extension.

Pools of forward and reverse probes were conjugated with both forward and reverse primers separately by mixing 22.5 μM primer with 10 μM total probe concentration, in 0.6x PBS. Each mixture was incubated at 60 °C for 16 hours. After incubation, the conjugates were purified using a modified Agencourt AMPure XP Kit (Beckman Coulter) and eluted in 20 μL 0.1x IDTE (IDT). A 2:1 bead to sample ratio was used according to the manufacturer’s instructions, except that prior to use, the bead buffer was extracted and replaced with an equal volume of a custom formulated buffer. The custom buffer consisted of 30% w/v PEG-8000, 1 M NaCl, 0.05% v/v Tween 20, 10 mM Tris-HCl, and 1 mM EDTA (all reagents from Sigma-Aldrich). Following conjugation, PDPs were quantified using the Qubit ssDNA Assay (ThermoFisher Scientific). Conjugates were made and then stored at −20 °C. PDPs consisting of forward probes with forward primers were labelled as FF, reverse probes with forward primers RF, and so on for all four combinations.

### Sample Sources

Four sample types were used in this study: mechanically sheared cell line DNA, enzymatically sheared cell line DNA, plasma-derived cell-free DNA (cfDNA), and FFPE-derived DNA. Mechanically sheared DNA was obtained from Horizon Discovery in mutant (HD786) and wild-type (HD776) standards (Samples 1 and 3, respectively, from Table 2). Mechanical shearing was performed by the manufacturer such that around 60% of the templates were within 100 bp to 250 bp, with fragments up to 400 bp. Mutation levels were measured by the manufacturer using droplet digital PCR. Enzymatically sheared cell line was generated from genomic DNA (HD753, Horizon Discovery), following the protocol described below. cfDNA was isolated from single donor human plasma samples (IPLAS – K2 EDTA, Innovative Research), as described below. FFPE-derived-DNA was obtained from Horizon Discovery, part number HD799.

### Cell-free DNA Extraction

First, 5 mL of plasma was centrifuged for 10 min at 2,000g. cfDNA was isolated from each sample using the QIAamp Circulating Nucleic Acid Kit (Qiagen) according to the manufacturer’s instructions. DNA was eluted from the column in 0.1x IDTE in a two-step process to maximize elution yield: 50 μL of 0.1x IDTE was incubated in the column for 10 min, followed by a 20,000g spin for 3 min; the column was then re-eluted after a 3 min incubation with another 50 μL 0.1x IDTE for a total elution volume of 100 μL. The DNA sample was further purified to remove any potential inhibitors using the Agencourt AmPure XP Kit (Beckman Coulter). A 1.4:1 bead to sample volumetric ratio was used as per manufacturer’s instructions, with the sample eluted in 0.1x IDTE. Extracted and purified DNA was then used directly for library preparation, or in cases where library preparation did not proceed within 24 hours, was frozen at −20 °C.

Following DNA extraction, sample concentration was measured using the Qubit dsDNA HS kit (ThermoFisher Scientific) as per the manufacturer’s instructions and used to calculate the number of human genome equivalent copies in each sample.

### FFPE DNA Pre-treatment

FFPE-derived DNA was pre-treated to reduce the impact of potential DNA damage, before target capture. 100 ng of DNA, as quantified by the Qubit dsDNA HS kit (ThermoFisher Scientific), was digested with 1 unit of UDG enzyme (New England Biolabs (NEB)) in a 50 μL reaction in 1X of the supplied reaction buffer (NEB). The mixture was incubated at 37 °C for 10 minutes, cooled to 4 °C, and immediately purified with the Agencourt AmPure XP Kit at a 3:1 bead to sample volumetric ratio as per manufacturer’s instructions. Samples were eluted in 20 μL of 10 mM Tris-HCl, pH 8. Total amplifiable DNA was quantified using KAPA hgDNA Quantification and QC Kit (KAPA Biosystems) as per the manufacturer’s instructions.

### Enzymatic DNA Shearing

Prior to enzymatic shearing, a buffer exchange was performed with cell line and FFPE-derived DNA samples with the Agencourt AmPure XP Kit at a 3:1 bead to sample volumetric ratio according to the manufacturer’s instructions. Samples were eluted in 40 μL of 10 mM Tris-HCl, pH 8. Cell line and FFPE-derived DNA samples were then enzymatically sheared immediately before ligation using the KAPA HyperPlus kit (KAPA Biosystems) according to the manufacturer’s instructions. 50 ng of cell line DNA or 40 ng of FFPE-derived DNA in 35 μL volume was added to 10 μL KAPA fragmentase (KAPA Biosystems) and topped up to a final volume of 50 μL in 1x supplied reaction buffer. Samples were incubated at 37 °C for 30 minutes, afterwards proceeding immediately to the A-tailing step of adapter ligation (described below). Shearing conditions were chosen as per manufacturer’s instructions to achieve a mode fragment length of 150 bp.

### Adapter Ligation

The KAPA Hyper Prep kit (KAPA Biosystems) was used as per the manufacturer’s instructions, with a 15 minute ligation incubation and a 200:1 adapter to insert ratio. Custom ligation adapter sequences were ordered for the LTC workflow (IDT), consisting of AGCACGCACCGAGATCTACAC**BBBB**ACACTCTTTCCCTACACGACGCTCTTCC GATCTT annealed to AGATCGGAAGAGCACACGTCTGAACTCCAGTCAC **BBBBNNNN** ACCGATCTCGTAACTCAGCGG, where BBBB indicates a four base sample-specific barcode for multiplexing samples on the sequencer, and NNNN indicates a four base UMI. The UMI-containing adapter was phosphorylated on its 5’ and 3’ ends. The non-UMI adapter contained a phosphorothioate bond between the last two bases on the 3’ end of the adapter. After ligation, the ligation mixture was purified using the Agencourt AMPure XP Kit (Beckman Coulter) as per manufacturer’s specification, with a 0.4:1 bead to sample volumetric ratio, and eluted in 40 μL of 0.1x IDTE. After elution, the sample was topped up to 100 μL with 0.1x IDTE. An additional cleanup with the Zymo Select-a-Size DNA Clean & Concentrator column (Zymo Research) was performed, as per the manufacturer’s instructions. A 5:1 binding buffer to ethanol ratio was used to select the desired product size. The final product was eluted in 25 μL of 0.1x IDTE. After cleanup, the entire volume of ligated DNA was amplified with custom primers TTTTTAGCACGCACCGAGATCTACAC and TTTTTCCGCTGAGTTACGAGATCGGT. Amplification proceeded for eight cycles with 0.3 μM of each primer, in 1x KAPA HiFi HotStart ReadyMix (KAPA Biosystems). Annealing was performed at 60 °C for 30 s, extension at 72 °C for 20 s, and denaturing at 98 °C for 20 s. The amplified products were cleaned up using the Agencourt AMPure XP Kit as per manufacturer’s specification, with a 1.2:1 bead to sample volumetric ratio, and eluted in 20 μL of 0.1x IDTE. The cleaned up template DNA was then quantified using the Qubit dsDNA HS kit (ThermoFisher Scientific) as per the manufacturer’s instructions.

### Target Capture

Target-capture-PCR (tcPCR) for the 35-gene panel was performed in two subsequent steps, each consisting of two reactions per sample. In the first step, the PDPs with untailed primers were used, split into two 50 μL reactions such that in the first reaction FF and RR PDPs were used to capture one sense of the target, and in the second reaction FR and RF PDPs were used for the other sense. Each reaction consisted of 5 nM of each individual PDP, 15 ng template DNA, 5 units of Platinum Taq polymerase (ThermoFisher Scientific), 4 mM MgCl_2_, 0.2 mM dNTP (Invitrogen) in 1x Platinum Taq Buffer (ThermoFisher Scientific). 15 tcPCR cycles were performed with a 30 s denaturing step at 95 °C followed by a combined annealing and extension step at 66 °C for 105 s. The ramp rate was 4 °C/s between 95 °C and 85 °C, and then 0.2 °C/s from 85 °C to 66 °C. The second tcPCR was performed using 12.5 uL of the amplified material from the first tcPCR, and was otherwise identical to the first step, with the following exceptions: PDPs with the tailed primers were used, ramp rate was 4 °C/s throughout cycling, 12 cycles were performed, and the combined annealing and extension step was done at 68 °C. Following amplification, libraries were purified using two back-to-back bead cleanups, using the Agencourt AMPure XP Kit as per manufacturer’s specification, with a 0.8:1 bead to sample volumetric ratio. Final libraries were eluted in 20 μL of 0.1x IDTE. tcPCR for the 4-gene panel was performed using a similar but earlier version of the protocol, that was the same with the exception of the following differences: reaction volume was 25 μL for both tcPCRs, 20 cycles were used in the first tcPCR, and 18 in the second, 6.25 μL of the first tcPCR was carried over into the second, the second tcPCR was eluted in 15 μL after cleanup.

### Sequencing and Data Analysis

Targeted libraries were sequenced on an Illumina MiSeq or MiniSeq with paired-end 2 x 150 bp reads as per manufacturer’s instructions. Prior to sequencing, samples were quantified using the KAPA Library Quant Kit (KAPA Biosystems) as per manufacturer’s instructions. Resulting FASTQ files were demultiplexed by sample barcode using Fulcrum Genomic’s FGBIO open source bioinformatics tool suite (https://github.com/fulcrumgenomics/fgbio) and then adapter-trimmed using Trimmomatic V0.36 (30). Trimmed read pairs were combined and aligned to the GRCh38/hg38 reference sequence using BWA-MEM (https://github.com/lh3/bwa) and output in BAM format. SAMtools (31) was then used for sorting and indexing BAM files. The resulting BAM files were grouped into UMI consensus reads by FGBIO for low level SNV detection. Picard Tools 2.9 (https://github.com/broadinstitute/picard) was then used to collect hybrid selection metrics, including on-target fraction, mean coverage and insert length distributions. SNV, CNV and indel mutation calling was achieved using GATK4 (https://software.broadinstitute.org/gatk/gatk4). CNV detection was not quantified, but CNVs were identified as “amplified”, “deleted” or “copy-number neutral” by the GATK4 CallCopyRatioSegments caller. Fusion detection was measured by comparing Picard de-duplicated reads containing alignments to both the CCDC6 and RET genes. Analysis outputs for assay validation and variant detection can be found in Supplementary Material S2 and S3, respectively.

## Acknowledgments

The authors wish to thank Integrated DNA Technologies Inc. (IDT) for support in probe synthesis. Additionally, the authors wish to acknowledge David Broemeling for his help in creating the figures, and Matthew Wiggin for his helpful review and discussions of the manuscript.

## Author Contributions

Conceived and designed the experiments: JP, AL, LG, MD, WC, LU, AM

Performed the experiments: AL, LG, MD, WC, LU

Analyzed the data: JP, AL, LG, MD, WC, LU, GS

Wrote the paper: JP

Edited the paper: AM

## References

1. Mamanova L, Coffey AJ, Scott CE, Kozarewa I, Turner EH, Kumar A, et al. Target-enrichment strategies for next-generation sequencing. Nat Methods. 2010;7(2):111–8.

2. Beadling C, Neff TL, Heinrich MC, Rhodes K, Thornton M, Leamon J, et al. Combining highly multiplexed PCR with semiconductor-based sequencing for rapid cancer genotyping. J Mol Diagn. 2013;15(2):171–6.

3. Tewhey R, Warner JB, Nakano M, Libby B, Medkova M, David PH, et al. Microdroplet-based PCR enrichment for large-scale targeted sequencing. Nat Biotechnol. 2009;27(11):1025–31.

4. Schenk D, Song G, Ke Y, Wang Z. Amplification of overlapping DNA amplicons in a single-tube multiplex PCR for targeted next-generation sequencing of BRCA1 and BRCA2. PLoS One. 2017;12(7):e0181062.

5. Satterfield BC. Cooperative primers: 2.5 million-fold improvement in the reduction of nonspecific amplification. J Mol Diagn.2014. p. 163–73.

6. Gentalen E, Chee M. A novel method for determining linkage between DNA sequences: hybridization to paired probe arrays. Nucleic Acids Res. 1999;27(6):1485–91.

7. Ståhlberg A, Krzyzanowski PM, Jackson JB, Egyud M, Stein L, Godfrey TE. Simple, multiplexed, PCR-based barcoding of DNA enables sensitive mutation detection in liquid biopsies using sequencing. Nucleic Acids Res. 2016;44(11):e105.

8. Zheng Z, Liebers M, Zhelyazkova B, Cao Y, Panditi D, Lynch KD, et al. Anchored multiplex PCR for targeted next-generation sequencing. Nat Med. 2014;20(12):1479–84.

9. Dahl F, Gullberg M, Stenberg J, Landegren U, Nilsson M. Multiplex amplification enabled by selective circularization of large sets of genomic DNA fragments. Nucleic Acids Res. 2005;33(8):e71.

10. Porreca GJ, Zhang K, Li JB, Xie B, Austin D, Vassallo SL, et al. Multiplex amplification of large sets of human exons. Nat Methods. 2007;4(11):931–6.

11. Hopmans ES, Natsoulis G, Bell JM, Grimes SM, Sieh W, Ji HP. A programmable method for massively parallel targeted sequencing. Nucleic Acids Res. 2014;42(10):e88.

12. Gunderson KL, Steemers FJ, Lee G, Mendoza LG, Chee MS. A genome-wide scalable SNP genotyping assay using microarray technology. Nat Genet. 2005;37(5):549–54.

13. Gnirke A, Melnikov A, Maguire J, Rogov P, LeProust EM, Brockman W, et al. Solution hybrid selection with ultra-long oligonucleotides for massively parallel targeted sequencing. Nat Biotechnol. 2009;27(2):182–9.

14. Albert TJ, Molla MN, Muzny DM, Nazareth L, Wheeler D, Song X, et al. Direct selection of human genomic loci by microarray hybridization. Nat Methods. 2007;4(11):903–5.

15. Bodi K, Perera AG, Adams PS, Bintzler D, Dewar K, Grove DS, et al. Comparison of commercially available target enrichment methods for next-generation sequencing. J Biomol Tech. 2013;24(2):73–86.

16. Schmitt MW, Fox EJ, Prindle MJ, Reid-Bayliss KS, True LD, Radich JP, et al. Sequencing small genomic targets with high efficiency and extreme accuracy. Nat Methods. 2015;12(5):423–5.

17. García-García G, Baux D, Faugère V, Moclyn M, Koenig M, Claustres M, et al. Assessment of the latest NGS enrichment capture methods in clinical context. Sci Rep. 2016;6:20948.

18. Alcaide M, Yu S, Davidson J, Albuquerque M, Bushell K, Fornika D, et al. Targeted error-suppressed quantification of circulating tumor DNA using semi-degenerate barcoded adapters and biotinylated baits. Sci Rep. 2017;7(1):10574.

19. Lee C, Bae JS, Ryu GH, Kim NKD, Park D, Chung J, et al. A Method to Evaluate the Quality of Clinical Gene-Panel Sequencing Data for Single-Nucleotide Variant Detection. J Mol Diagn. 2017;19(5):651–8.

20. Underhill HR, Kitzman JO, Hellwig S, Welker NC, Daza R, Baker DN, et al. Fragment Length of Circulating Tumor DNA. PLoS Genet. 2016;12(7):e1006162.

21. Newman AM, Bratman SV, To J, Wynne JF, Eclov NC, Modlin LA, et al. An ultrasensitive method for quantitating circulating tumor DNA with broad patient coverage. Nat Med. 2014;20(5):548–54.

22. Ware JS, John S, Roberts AM, Buchan R, Gong S, Peters NS, et al. Next generation diagnostics in inherited arrhythmia syndromes: a comparison of two approaches. J Cardiovasc Transl Res. 2013;6(1):94–103.

23. Wang Q, Wang X, Tang PS, O’leary GM, Zhang M. Targeted sequencing of both DNA strands barcoded and captured individually by RNA probes to identify genome-wide ultra-rare mutations. Sci Rep. 2017;7(1):3356.

24. Caruccio N. Preparation of next-generation sequencing libraries using Nextera(tm) technology: simultaneous DNA fragmentation and adaptor tagging by in vitro transposition. Methods Mol Biol. 2011;733:241–55.

25. Snyder MW, Kircher M, Hill AJ, Daza RM, Shendure J. Cell-free DNA Comprises an In Vivo Nucleosome Footprint that Informs Its Tissues-Of-Origin. Cell. 2016;164(1-2):57–68.

26. Mamanova L, Turner DJ. Low-bias, strand-specific transcriptome Illumina sequencing by on-flowcell reverse transcription (FRT-seq). Nat Protoc. 2011;6(11):1736–47.

27. Newman AM, Lovejoy AF, Klass DM, Kurtz DM, Chabon JJ, Scherer F, et al. Integrated digital error suppression for improved detection of circulating tumor DNA. Nat Biotechnol. 2016;34(5):547–55.

28. Tewhey R, Nakano M, Wang X, Pabón-Peña C, Novak B, Giuffre A, et al. Enrichment of sequencing targets from the human genome by solution hybridization. Genome Biol. 2009;10(10):R116.

29. Dwight Z, Palais R, Wittwer CT. uMELT: prediction of high-resolution melting curves and dynamic melting profiles of PCR products in a rich web application. Bioinformatics. 2011;27(7):1019–20.

30. Bolger AM, Lohse M, Usadel B. Trimmomatic: a flexible trimmer for Illumina sequence data. Bioinformatics. 2014;30(15):2114–20.

31. Li H, Handsaker B, Wysoker A, Fennell T, Ruan J, Homer N, et al. The Sequence Alignment/Map format and SAMtools. Bioinformatics. 2009;25(16):2078–9.

